# Development of a simple *in vitro* assay to identify and evaluate nucleotide analogs against SARS-CoV-2 RNA-dependent RNA polymerase

**DOI:** 10.1101/2020.07.16.205799

**Authors:** Gaofei Lu, Xi Zhang, Weinan Zheng, Jialei Sun, Lan Hua, Lan Xu, Xin-jie Chu, Sheng Ding, Wen Xiong

## Abstract

Nucleotide analogs targeting viral RNA polymerase have been approved to be an effective strategy for antiviral treatment and are attracting antiviral drugs to combat the current SARS-CoV-2 pandemic. In this report, we develop a robust *in vitro* nonradioactive primer extension assay to evaluate the incorporation efficiency of nucleotide analog by SARS-CoV-2 RNA-dependent RNA polymerase (RdRp) quantitively. Our results show that many nucleotide analogs can be incorporated into RNA by SARS-CoV-2 RdRp, and that the incorporation of some of them leads to chain termination. The discrimination values of nucleotide analog over those of natural nucleotide were measured to evaluate the incorporation efficiency of nucleotide analog by RdRp. We found that the incorporation efficiency of Remdesivir-TP is higher than ATP, and we did not observe chain termination or delayed chain termination caused by single Remdesivir-TP incorporation, while multiple incorporations of Remdesivir-TP caused chain termination in our assay condition. The incorporation efficiency of Ribavirin-TP and Favipiravir-TP is very low either as ATP or GTP analogs, which suggested that mutagenesis may not be the mechanism of action of those two drugs against SARS-CoV-2. Incorporation of Sofosbuvir-TP is also very low suggesting that sofosbuvir may not be very effective in treating SARS-CoV-2 infection. As a comparison, 2’-C-Methyl-GTP can be incorporated into RNA efficiently, and the derivative of 2’-C-Methyl-GTP may have therapeutic application in treating SARS-CoV-2 infection. This report provides a simple screening method that should be useful in evaluating nucleotide-based drugs targeting SARS-CoV-2 RdRp, and for studying the mechanism of action of selected nucleotide analog.

## Introduction

An ongoing global pandemic of Coronavirus disease 2019 (COVID-19) was caused by severe acute respiratory syndrome coronavirus 2 (SARS-CoV-2) (1–3), which is single-stranded, positive-sense RNA virus and has the largest genomes (around 30kb) known among RNA virus (4). Right now, SARS-CoV-2 is continuing its spread across the world, and currently, no effective vaccines are available, thus the search for safe and effective small-molecule inhibitors that would be active against SARS-CoV-2 represents a high research priority.

Nucleoside/nucleotide-based inhibitors targeting viral DNA or RNA polymerase have been proven to be the backbone of antiviral therapies (5, 6). Currently, there are over 25 approved therapeutic nucleosides/nucleotides used for the therapy of various viral infections of high medical importance, such as HIV (tenofovir, AZT, et al), hepatitis B (lamivudine), hepatitis C (sofosbuvir), or herpes infection (acyclovir). The proven success in using nucleoside-based drugs to effectively treat viral infections and the current knowledge of the development pathway for developing this class of inhibitors makes them attractive antiviral agents to combat the current pandemic of COVID-19 (7–9).

Following the intracellular phosphorylation, nucleotide analog 5’-triphosphates (the active form of nucleoside analogs) are competitively incorporated into nascent viral RNA chains and thus inhibit viral replication (5). The mechanism of action varies for different nucleotide analogs. The most common mechanism of action is chain termination caused by the incorporation of nucleotide analogs resulting in the formation of incomplete viral RNA chains (10, 11). Non-chain terminators may also be used as antiviral drugs, such as Ribavirin and Favipiravir which could induce mutagenesis due to their ambiguous base-pairing properties after incorporated into viral RNA (12–17).

Like other Coronaviruses, SARS-CoV-2 encodes an RNA-dependent-RNA polymerase (RdRp) on the nsp12 gene (4). This protein, which catalyzes RNA synthesis, forms a replication complex with nsp7, nsp8, and other virally encoded proteins and host proteins that are responsible for mRNA synthesis, as well as the synthesis of genomic RNA for progeny viruses (8). The structure of the nsp12-nsp8-nsp7 complex was recently determined using cryo-EM (18). Additionally, unlike other viruses, SARS-CoV-2 encodes an exonuclease gene (nsp14) and can perform proofreading activity to remove the mismatched nucleotides from viral RNA (19, 20). This probably is one of the difficulties to develop nucleotide polymerase inhibitors against SARS-CoV-2 and this proofreading activity of SARS-CoV-2 should be taken into consideration when developing a nucleoside-based antiviral drug against SARS-CoV-2. Nucleotide analogs with different mechanisms of action (chain terminator versus non-chain terminator) may respond differently to SARS-CoV-2 proofreading activity mediated by nsp14. So far, numerous nucleoside/nucleotide analogs have been described to inhibit SARS-CoV-2 including Remdesivir, Ribavirin, BCX4430, Gemcitabine hydrochloride, β-D-N4-hydroxycytidine and 6-Azauridine (7). The detailed mechanism of action of those nucleotide analogs against SARS-CoV-2 still needs to be addressed.

In this report, we developed a robust *in vitro* nonradioactive primer extension assay using a fluorescently labeled RNA primer annealed to an RNA template. The incorporation efficiency and chain termination ability of a series of nucleotide analogs were determined using this assay. The method is very valuable for evaluation of the antiviral potential of nucleotide analogs and the structure-activity information derived from these studies can be used to explore the mechanism of action of given nucleotide inhibitors, and design nucleotide analogs with better properties.

## Materials and methods

### Chemicals

ATP, UTP, CTP, GTP were purchased as 100 mM solution from Thermo Fisher Scientific (Massachusetts, USA). Urea, taurine, DTT, MgCl_2_, imidazole, IPTG were purchased from Bidepharm (Shanghai, China). LB medium, NaCl, HisPur Ni-nitrilotriacetic acid agarose resin were purchased from Thermo Fisher Scientific (Massachusetts, USA). Fluorescently labeled RNA oligonucleotides, as well as unlabeled RNA oligonucleotides, were chemically synthesized and purified by HPLC by Genscript (Nanjing, China). Stellar competent cells were purchased from Takara (Mountain, California, USA). BL21 (DE3) cells were purchased from TransGen Biotech (Beijing, China). The pMal-c5X vector was purchased from New England Bioscience (NEB, Ipswich, MA). Tenofovir-DP, 2’-C-Methyl-CTP, and 2’-C-Methyl-GTP were purchased from Carbosynth (San Diego, CA). 6-Chloropurine-TP, Gemcitabine-TP, 6-Methyl-thio-GTP, Clofarabine-TP, 8-Oxo-GTP, 6-Thio-GTP, Stavudine-TP, and Lamivudine-TP were purchased from Jena Bioscience (Jena, Germany). Ribavirin-TP was purchased from Santa Cruz Biotechnology (Texas, USA). Favipiravir-TP was purchase from Toronto Research Chemicals (TRC, Toronto, Canada). 2’-Amino-UTP, 2’-O-Methyl-UTP, ara-UTP, and 2’-Azido-UTP were purchased from Trilink BioTechnologies (San Diego, USA). Sofosbuvir-TP was purchased from MedChemExpress (MCE, New Jersey, USA). Remdesivir-TP was custom synthesized in Wuxi AppTech (Shanghai, China).

### Nsp12 protein expression and purification

SARS-CoV-2 nsp12 gene (amino acid 4393-5324 Uniprot: P0DTD1) was synthesized de novo by GenScript (Nanjing, China) and constructed onto pET22b vector between *NdeI* and *XhoI* sites. Nsp12 was expressed with a C-terminal 10×His-tag in BL21 (DE3) cells at 16°C; 2 mM MgCl_2_ and 50 μM ZnSO_4_ were supplemented in culture during induction. After overnight cultivation, cells were harvested and lysed by high-pressure homogenizer in buffer containing 25 mM HEPES pH 7.5, 150 mM NaCl, 4 mM MgCl_2_, 50 μM ZnSO_4_, 10% glycerol, 2.5 mM DTT and 20 mM imidazole. Cell debris was removed by centrifugation at 13 000 rpm. Nsp12 was then purified by Nickel-affinity chromatography followed by ion-exchange chromatography (using HisTrap FF and Capto HiResQ 5/50 column, respectively. GE Healthcare, USA). Nsp12 eluates within conductivity of 19-23 mS/cm were combined and injected onto a Superdex 200 increase 10/300 GL column (GE Healthcare, USA) in a buffer containing 25 mM HEPES pH 7.5, 250 mM NaCl, 1 mM MgCl_2_, 1 mM Tris(2-carboxyethyl)phosphine and 10% glycerol. Peak fractions were combined, concentrated to 10 μM and stored at −80°C before enzymatic assay. Nsp12 loss-of-function mutant (SDD→SAA mutation, amino acid 5151-5153, Uniprot: P0DTD1) was prepared in an identical process. Protein identification by Mass spectrometry was performed by Tsinghua University.

### Nsp7 and nsp8 protein expression and purification

SARS-CoV-2 nsp8 gene (nucleotide 12092-12685, strain, GenBank: MN908947.3) was synthesized de novo by GenScript (Nanjing, China) and cloned into a pMal-c5X vector under tac promoter control (without a maltose-binding protein sequence). SARS-CoV-2 nsp7 gene (nucleotide 11846-12091, GenBank: MN908947.3) was synthesized de novo by GenScript (Nanjing, China) and cloned into the pMal-c5X vector using the same way as nsp8. The expression plasmids were transformed into Stellar competent cells. Protein expression was induced at 16 °C overnight night by addition of 0.3 mM IPTG. Cells were harvested and cell pellets were resuspended in cell lysis buffer (20 mM HEPES, pH 7.5, 10% glycerol, 100 mM NaCl, 0.05% Tween 20, 10 mM DTT, 1 mM MgCl_2_, 20 mM imidazole, 1 × protease inhibitor cocktail). Cell disruption was performed at 4 °C for 10 min using a high-pressure homogenizer. The cell extract was clarified by centrifugation at 12000 RPM for 10 min at 4 °C. Nsp7 and nsp8 were purified by HisPur Ni-NTA agarose resin and the enzymes were eluted from the resin with elution buffer (20 mM HEPES, pH 7.5, 50 mM NaCl, 300 mM imidazole, 10 mM DTT, 0.01% Tween 20). The eluted enzymes were adjusted to 40% glycerol and store at −80 °C. Protein identification by Mass spectrometry was performed by Tsinghua University.

### Primer and template annealing

To generate RNA primer-template complexes, 1 μM fluorescently (Cy5.5) labeled RNA primer and 5 μM unlabeled RNA template were mixed in 50 mM NaCl in deionized water, incubated at 98 °C for 10 min, and then slowly cooled to room temperature. The annealed primer and template complexes (P/Ts) were stored at −20 °C before use in primer extension assay.

### Primer extension assay

The ability of RNA synthesis by purified polymerase was determined in a primer extension reaction using P/Ts prepared by annealing Cy5.5 labeled RNA primer and unlabeled RNA template (described above). A typical primer extension reaction was performed in a 10 μl reaction mixture containing reaction buffer (20 mM HEPES, pH 7.5, 5 mM MgCl_2_, 10 mM DTT, 0.01% Tween 20), 5 nM P/T, and 50 nM nsp12, 2 μM nsp8-7 (copurified, contain 2 μM nsp8 and 10 μM nsp7) unless otherwise specified.. The reaction was initiated by the addition of rNTPs at a final concentration of 100 μM, unless otherwise specified, followed by incubation for 1 hour at 37 °C. The reactions were quenched by the addition of 20 μl stopping solution (8 M urea, 90 mM Tris base, 29 mM Taurine, 10 mM EDTA, 0.02% SDS, 0.1% bromophenol blue). The quenched samples were denatured at 95 °C for 10 min, and the primer extension products were separated using 10% denaturing polyacrylamide gel electrophoresis (urea-PAGE) in 1 × TTE buffer (90 mM Tris base, 29 mM taurine, 0.5 mM EDTA). After electrophoresis, the gels were scanned using an Odyssey infrared imaging system (LI-COR Biosciences, Lincoln, NE). The images were analyzed, and the proper RNA bands were quantified using Image Studio Lite (version 5.2; LI-COR Bioscience, Lincoln, NE). Data were analyzed using GraphPad Prism 7.

### Analysis of chain termination ability of nucleotide analogs

The primer extension reactions were performed as described above. Incorporation and chain termination of tested nucleotides were measured in two separate assays. Nucleotide analog incorporation assay: P/Ts (5 nM) and nsp12-nsp8-nsp7 complex (RdRp) (50 nM nsp12, 2 μM nsp8-7) were incubated with a natural rNTP (the first nucleotide to be incorporated) and tested nucleotide analogs (the second nucleotide to be incorporated) and the reactions were continued at 37 °C for 30 min before adding stopping solution. Chain termination assay: Incorporation of nucleotide analogs was performed as described above and after incorporation of nucleotide analogs, two natural rNTPs (third and fourth nucleotide to be incorporated) were added to the reaction mixture and reactions were continued at 37 °C for another 30 min before adding stopping solution. The quenched samples were heated at 95 °C for 10 min and analyzed by denaturing urea-PAGE as described above. The concentration and identity of nucleotides added for each reaction were described in the figure legend in Fig. 6.

### Measurement of nucleotide analog incorporation efficiency

Different P/Ts were designed to test individual analog using the same method described previously (21, 22). To perform the reaction, 5 nM P/T and 50 nM nsp12, 2 μM nsp8-nsp7 were incubated in reaction buffer in the presence of 0.1 μM the first natural ribonucleotide for 30 minutes at 37 °C, and then different concentrations of the nucleotide analogs to be tested were added to the reaction mixtures. The reactions were continued at 22 °C for the time indicated in each figure legend and subsequently quenched and analyzed by urea-PAGE as described above. After electrophoresis, the gels were scanned using the Odyssey infrared imaging system. The intensity of the different RNA bands was quantified using Image Studio Lite. The incorporation efficiencies of the different nucleotide analogs were evaluated by measurement of the *K*_1/2_ values (the analog triphosphate concentrations resulting in 50% product extension) and the corresponding discrimination values (*D*_analog_, defined as *K*_1/2, analog_/*K*_1/2, natural nucleotide_ when both were measured under the same assay condition), as previously described (21, 22).

## Results

### Expression and purification of SARS-CoV-2 nsp12, nsp7, and nsp8

Numbers of reports have shown that functional SARS-CoV-2 nsp12-nsp8-nsp7 complex (RdRp) can be expressed and purified from insect cells (23). In this report, nsp12, nsp7, and nsp8 were successfully expressed and purified from *E. coli* (Fig. 1), and the functional nsp12-nap8-nsp7 complex was assembled by simply mixing nsp12, nsp7 and nsp8 together *in vitro*. Nsp12 was expressed as C-terminal His-tagged protein and purified according to the published papers with modification (18, 24). Nsp7 and nsp8 were expressed as N-terminal His-tagged protein and protein purification was performed using Nickel agarose resin. It was reported that pre-formed nsp7 and nsp8 complex has better ability to promote nsp12 polymerase activity (25), copurification of nsp7 and nsp8 was also performed by simply mixing cells expressing nsp8 and nsp7 before lysing cells, which will allow nsp7 and nsp8 to form a stable complex during purification. Nsp8-7 was used to represent copurified nsp8 and nsp7 in this report. After purification, the identity of the purified proteins was confirmed by mass spectrometry analysis (data not shown).

**Fig. 1.**
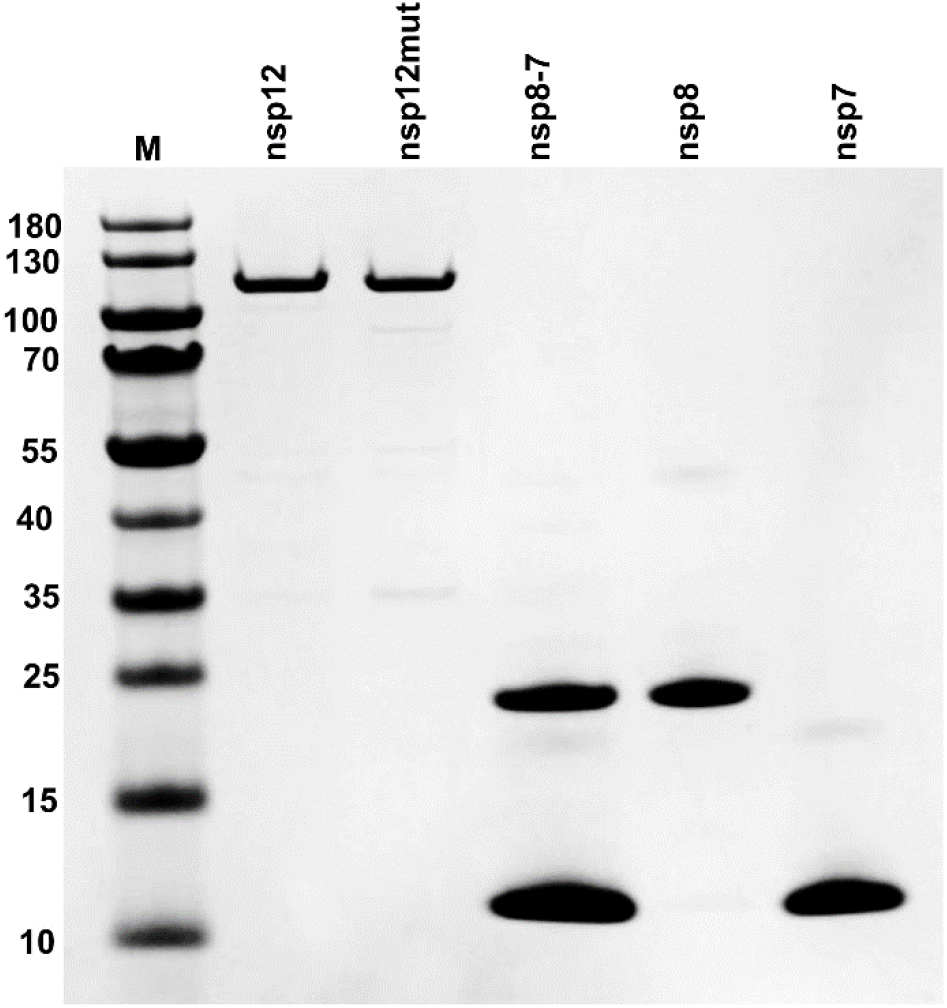
Expression and purification of nsp12, nsp7, nsp8, and nsp8-7. Nsp12mut contains an active site mutation (active site motif SDD→SAA). Nsp8-7 represents copurified nsp8 and nsp7. M, protein molecular weight markers. The sizes of protein markers (in kDa) are indicated on the left.

### Nsp12-nsp8-nsp7 complex assembles and activity measurement

To measure the polymerase activity of the purified enzyme, a primer extension assay employing an RNA template (40-mer RNA corresponding to the sequence of 3’-end of SARS-CoV-2 RNA genome) and a fluorescently labeled RNA primer (30-mer) was developed (Fig. 2A). The detail of the RNA primer extension method was described previously (21, 22, 26). It was reported that nsp7 and nsp8 could form a complex with nsp12, which is essential for RNA synthesis catalyzed by nsp12. In this experiment, different combinations of purified protein (nsp12, nsp7, nsp8, and nsp8-7) were used in the primer extension assay to test the RNA synthesis ability of different combinations (Fig. 2B). The result showed that nsp12, nsp7, or nsp8 alone did not have any measured RNA synthesis ability in the reaction condition described in this experiment (Fig. 2B, lane 2, 3, and 6). Nsp12 and nsp7 together also did not have RNA polymerase activity (Fig. 2B, lane 8), and nsp12 and nsp8 together had weak RNA polymerase activity (Fig. 2B, lane 9). Maximum polymerase activity was observed in the presence of nsp12, nsp8, and nsp7 together (Fig. 2B, lane 10, 11). An active-site mutation (SDD→SAA) of nsp12 (Nsp12mut) showed no RNA synthesis in the presence of nsp8 and nsp7, which confirm that RNA synthesis is mediated by nsp12. The ability of nsp8-7 to promote nsp12 polymerase was further evaluated through an enzyme dilution assay and compared with that of separately added nsp8 and nsp7 (Fig. 2C). The result showed that nsp8-7 had a better ability to promote nsp12 catalyzed RNA synthesis. Nsp8-7 was used in the following primer extension assays described in the report and nsp8 concentration in nsp8-7 was used to represent the concentration of nsp8-7.

**Fig 2.**
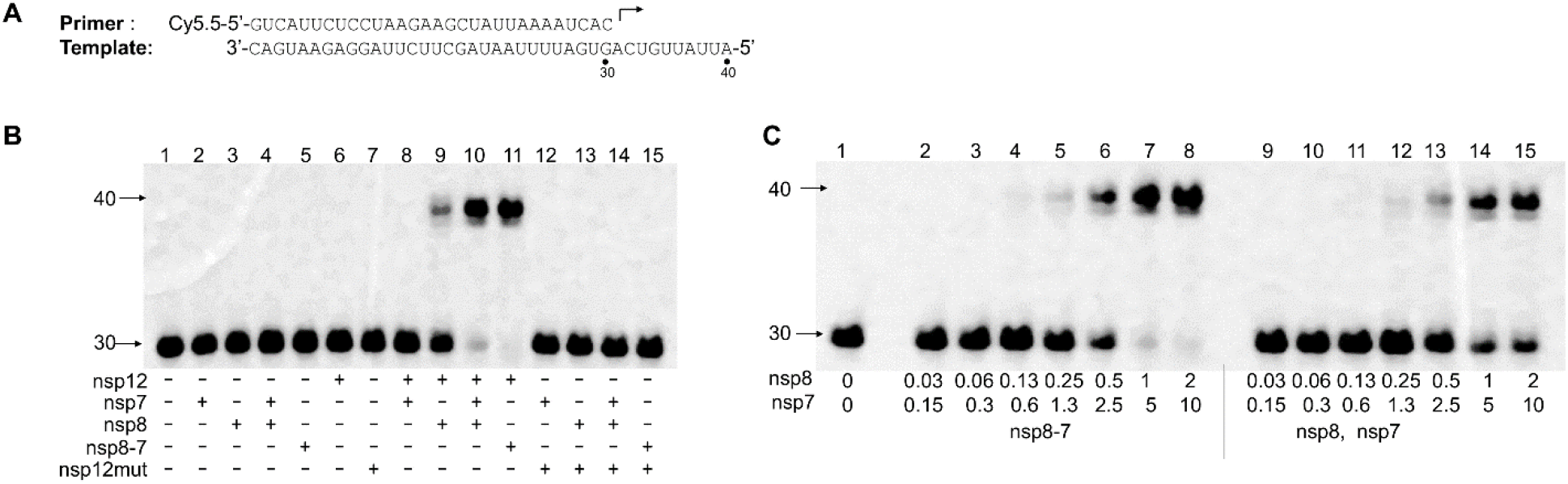
Analysis of nsp12 polymerase activity using a primer extension assay. **(A)** The RNA primer and template used in this assay. The 30-mer primer (top) contains a fluorescent label (Cy5.5) at the 5’ end and the arrow indicates the location and direction of primer extension to form a 40-mer product. **(B)** Analysis of the nsp12 polymerase activity in the presence of nsp7, nsp8 or nsp8-7 (co-purified nsp8 and nsp7). The enzymes used in each reaction are indicated at the bottom of the gel. The concentrations for nsp12, nsp12mut, nsp7, nsp8 and nsp8-7 (co-purified nsp8 and nsp7) are 50 nM, 50 nM, 10 μM, 2 μM, and 2 μM (2 μM nsp8, 10 μM nsp7), respectively. Different enzymes and P/T (5 nM) were incubated in reaction buffer and the reactions were initiated by the addition of 100 μM rNTPs, and then continued at 37 °C for 1 h, and then were stopped by adding stopping solution. The products were separated on denaturing polyacrylamide gels. **(C)** Primer extension activity using nsp12 and nsp8, nsp7 or nsp8-7 (copurified). Nsp12 (50 nM) was used in all the samples. Primer extension reaction was performed as described above. In lane 2-8, co-purified nsp8 and nsp7 was used, and in lane 9-15, nsp8 and nsp7 were added to the reaction separately. The final concentrations of nsp8, nsp7 in the reaction were labeled under each lane in micromolar.

### Primer extension assay development and optimization

It has been shown that the primer/template (P/T) scaffold could have a major influence on the efficiency of primer extension reaction catalyzed by mitochondrial DNA-dependent RNA polymerase (22). In this report, three RNA primers (Primer I, Primer II and Primer III) complementary to different regions of 3’-end of template (40-mer) were used to form three different P/T scaffolds (Fig.3A). The efficiency of RNA synthesis by SARS-CoV-2 RdRp using three P/Ts was tested in a primer extension assay with a serious dilution of nsp12 and a constant concentration of nsp8-7 (Fig. 3B). The result showed that the SARS-CoV-2 RdRp prefers longer primer (Primer III, 30mer) annealing with the RNA template. As a comparison, the efficiency of RNA synthesis using three P/Ts by dengue RdRp was similar (data not shown). Primer III (30-mer) shown in Fig. 3A was used in the following primer extension assays.

**Fig. 3.**
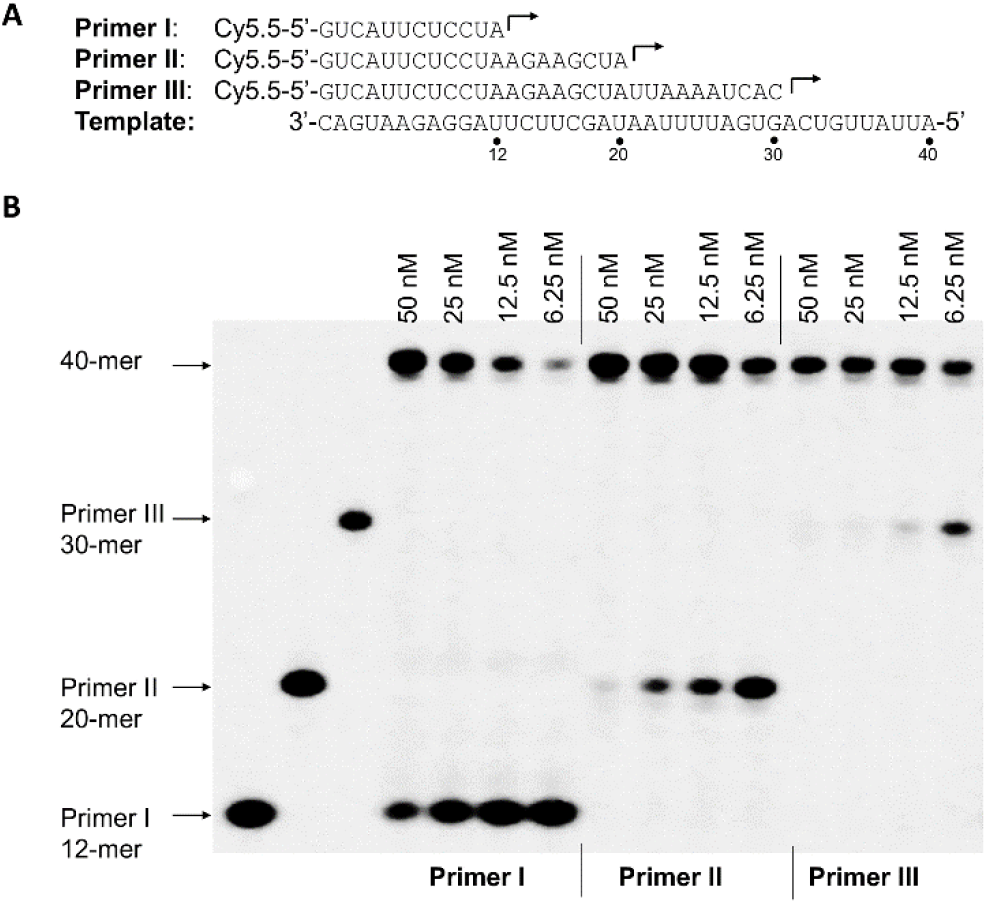
Influence of P/T scaffolds on the efficiency of the primer extension reaction catalyzed by SARS-CoV-2 RdRp. **(A)** The RNA primers (Primer I, Primer II and Primer III) and RNA template used in the experiments whose results are shown in panel B. **(B)** Primer extension assay using the different P/Ts from panel A. Four different nsp12 concentrations (6.25 nM, 12.5 nM, 25 nM, 50 nM; indicated on top of the gel), 2 μM nsp8-7 (2 μM nsp8 and 10 μM nsp7), and 5 nM different P/Ts (formed by annealing Primer I, Primer II or Primer III with the template shown in panel A; indicated at the bottom of the gel) were used in the assay. The reactions were initiated by the addition of 100 μM rNTP and continued at 37 °C for 1 h, and the products were separated on denaturing polyacrylamide gels. Three P/Ts without nsp12 added were used as a negative control (left three lanes) and the length of the primers (12-mer, 20-mer, 30-mer) was shown on the left of the gel.

To develop a robust polymerase assay, an optimal nsp8-nsp7 and nsp12 concentrations used in the reaction should be carefully determined. An enzyme dilution assay with different concentrations of nsp8-7 and nsp12 was performed (Fig. 4). The concentration of nsp8 was used to represent the concentration of co-purified nsp8-nsp7. The result showed that the primer extension products correlated with the concentration of nsp12 (Fig. 4C) and a higher concentration of nsp8-7 had better ability to promote nsp12 polymerase activity (Fig. 4D). In the following primer extension assays, 50 nM nsp12 and 2 μM nsp8-7 (2 μM nsp8 and 10 μM nsp7) were used. Under this assay condition, RNA primer (5nM) can be extended completely while consuming less nsp12 enzyme.

**Fig. 4.**
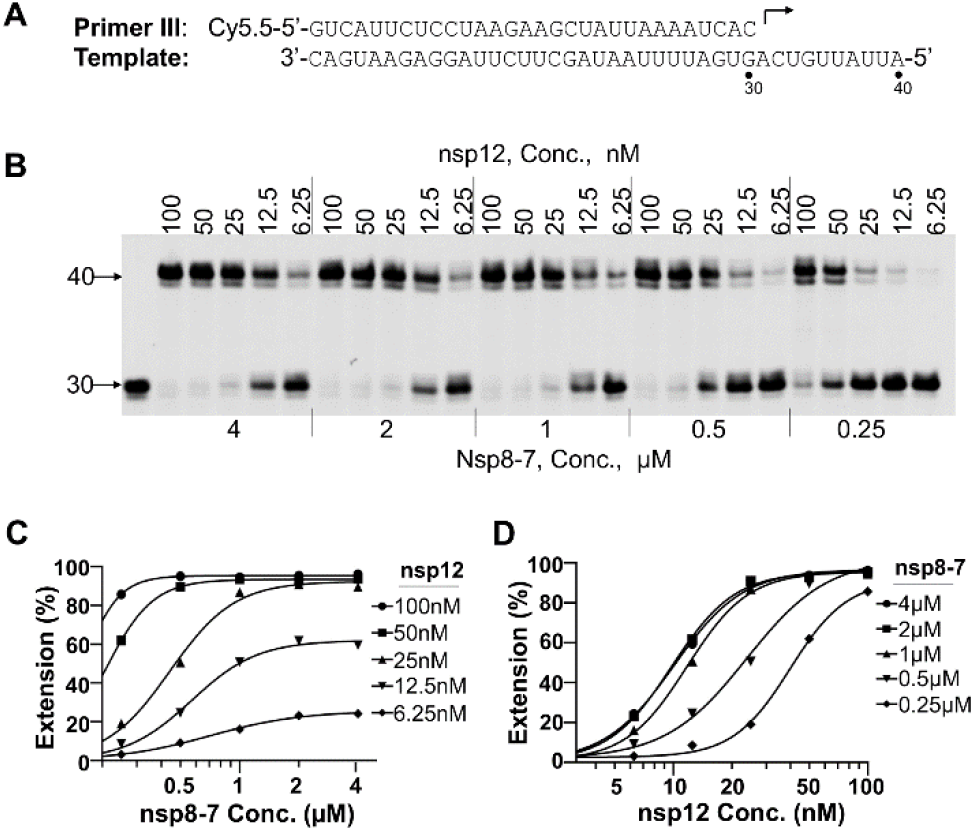
Nsp12 and nsp8-7 concentration optimization. **(A)** P/T used in this assay. **(B)** Analysis of the primer extension activity in the condition of different concentrations of nsp12 and nsp8-7. Serial dilutions of nsp12 (indicated on the top of the gel), serial dilutions of nsp8-7 (indicated at the bottom of the gel), and 5 nM P/T were used in this assay. The reaction was initiated by the addition of 100 μM rNTP, and the reactions were performed at 37 °C for 1 h and were then stopped by adding stopping solution. The products were separated on denaturing PAGE gels. The first lane on the left is the reaction without rNTP added which serves as a negative control. The number 30 and 40 on the left of the gels indicate the location of the 30-mer primer and 40-mer full-length product. **(C)** The percentage of full-length products in panel B was plotted against the nsp8-7 concentrations. The results were fitted to sigmoidal dose-response curves using the GraphPad Prism program. **(D)** The percentage of full-length products in panel B was plotted against the nsp12 concentrations. The results were fitted to sigmoidal dose-response curves using the GraphPad Prism program.

### Nucleotide incorporation and chain termination

To inhibit viral RNA synthesis or disrupt viral RNA function, a nucleotide analog must be incorporated into newly synthesized viral RNA by RNA polymerase. Several different nucleotide analogs (Fig. 5) that have been used as antiviral drugs or in the various applications were selected, and the utilization of those nucleotide analogs by SARS-CoV-2 RdRp and the ability of those analogs to cause chain termination (one of the major mechanisms of inhibition of viral RNA polymerase) after incorporation were evaluated in the primer extension assay (Fig. 6). Fig. 6 A and B show the incorporation and chain termination potential of several ATP analogs tested in the primer extension assay. A special template was designed for testing ATP analog as such that the first nucleotide to be incorporated is UTP and ATP or ATP analogs as the second nucleotide to be incorporated (Fig. 6A). The result showed that all the nucleotide analogs tested in this assay can be incorporated into RNA as ATP analogs at 100 μM (Fig. 6B. lane 3, 5, 7, 9, 11, 13, 15). Longer band in lane 5, 11, 13, 15 than the band in lane 3 shown in Fig. 6B suggested that 6-Chloropurine-TP, Remdesivir-TP, Ribavirin-TP, and Favipiravir-TP can also be incorporated as GTP (the third nucleotide to be incorporated) in this assay. After adding GTP and CTP (third and fourth nucleotide to be incorporated), if incorporation of a nucleotide analog can lead to chain termination, the band will not be extended (for example, comparing lane 9 and lane 10 in Fig. 6B), otherwise, it will be extended (for example, comparing lane 3 and lane 4 in Fig. 6B). The result showed that the incorporation of Tenofovir-DP caused chain termination (Fig. 6B, lane 10), and incorporation of 6-Choropurine-TP, Clofarabine-TP, Remdesivir-TP, Ribavirin-TP, and Favipiravir-TP did not cause chain termination (Fig. 6B, lane 6, 8, 12, 14, 16). In Fig. 6B, product in lane 6, 8, 12, 14, and 16 are shorter than the product shown in lane 4 (40-mer full-length product), which is probably because multiple ATP analog incorporations caused chain termination at various places.

**Fig. 5.**
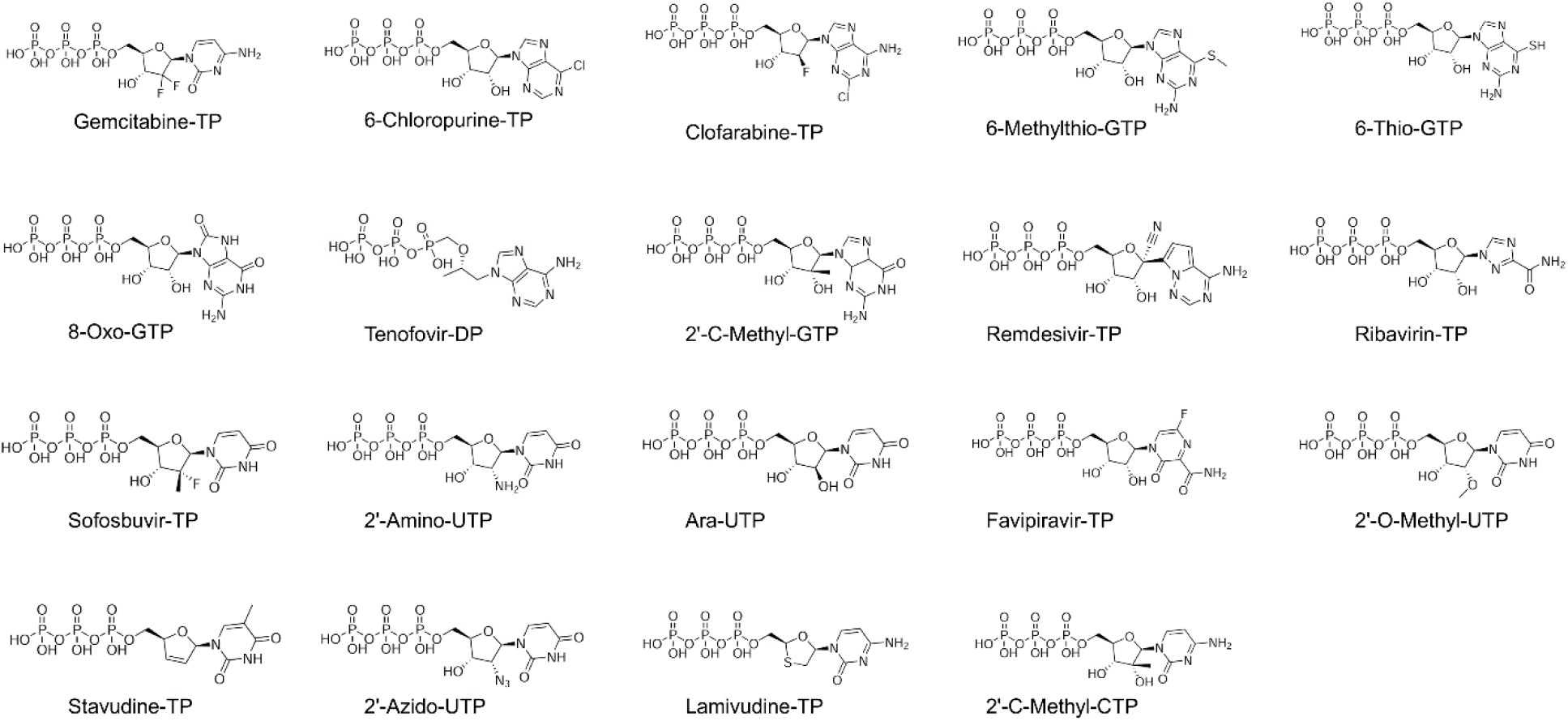
Chemical structure of the nucleotide 5’-triphosphate analogs tested in this study.

**Fig. 6.**
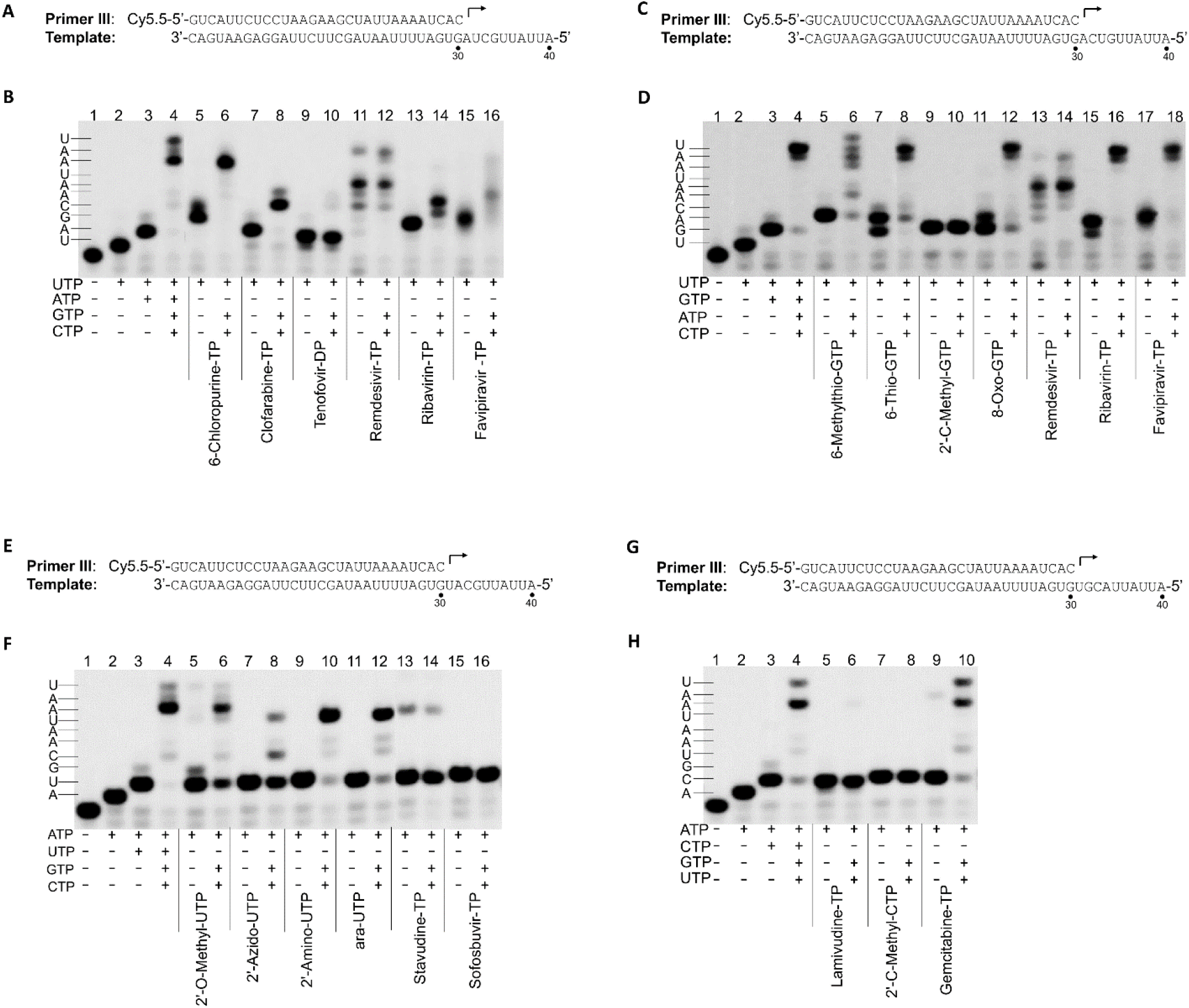
Analysis of incorporation and chain termination effects of nucleotide analogs in primer extension assay catalyzed by SARS-CoV-2 RdRp. **(A)** The P/T used in this assay whose result is shown in panel B. The first, second, third, and fourth nucleotides to be incorporated are UTP, ATP, GTP, and CTP, respectively. **(B)** Analysis of the incorporation and chain termination abilities of the ATP analogs. Incorporation and chain termination of tested nucleotides were measured in two separate assays. Incorporation (lane 3, 5, 7, 9, 11, 13, 15): primer extension reactions were initiated by the addition of 0.1 μM UTP and 100 μM ATP analogs, as indicated under each lane. The reactions were performed at 37 °C for 30 min and stopped by the addition of stopping solution. Chain termination ability (lane 4, 6, 8, 10, 12, 14, 16): primer extension reactions were initiated by the addition of 0.1 μM UTP and 100 μM ATP analogs, as indicated under each lane. The reactions were performed at 37 °C for 30 min, and then 0.1 μM GTP and 0.1 μM CTP were added to the reaction mixture and the reaction were continued at 37 °C for another 30 min before adding stopping solution. The reaction products were resolved by denaturing PAGE. Analysis of incorporation and termination of GTP analogs, UTP analogs, and CTP analogs were shown in panel **C-D**, panel **E-F** and panel **G-H** respectively using a similar method.

Fig.6C-D shows incorporation and chain termination potential of several GTP analogs. The result showed that all the analogs listed in Fig. 6D can be incorporated into RNA as GTP analog at 100 μM (Fig. 6D, lane 5, 7, 9, 11, 13, 15, and 17). Incorporation of 2’-C-Methyl-GTP caused immediate chain termination (Fig. 6D, lane 10); incorporation of 6-Methylthio-GTP, 6-Thio-GTP, 8-Oxo-GTP, Remdesivir-TP, Ribavirin-TP, and Favipiravir-TP did not cause chain termination (Fig. 6D, lane 6, 8, 12, 14, 16 and 18).

Fig. 6E-F shows incorporation and chain termination potential of several UTP analogs. The result showed all the analogs shown in Fig. 6F can be incorporated as UTP analogs at 100 μM (Fig. 6F, lane 5, 7, 9, 11, 13 and 15), incorporation of Sofosbuvir-TP and Stavudine-TP caused immediate chain termination (Fig. 6F, lane14, 16); incorporation of 2’-O-Methyl-UTP and 2’-Azido-UTP caused partial chain termination (Fig. 6F, lane 6 and 8); incorporation of 2’-Amino-UTP, and ara-UTP did not cause chain termination (Fig. 6F, lane 10 and 12).

Fig. 6G-H shows incorporation and chain termination potential of several CTP analogs. The result showed that all the analogs shown in Fig. 6H can be incorporated as CTP analogs at 100 μM (Fig. 6H, lane 5, 7 and 9), Lamivudine-TP and 2’-C-Methyl-CTP caused immediate chain termination (Fig. 6H, lane 6 and 8), and Gemcitabine-TP did not cause chain termination (Fig. 6H, lane 10).

### Incorporation efficiency of nucleotide analogs by RdRp

The relative incorporation efficiency of nucleotide analogs versus natural nucleotide (discrimination value) by viral RNA-dependent RNA polymerase has been used to evaluate the antiviral potential of nucleotide analogs targeting Zika virus and dengue virus RNA-dependent RNA polymerase (21), and it also has been used to evaluate potential mitochondrial toxicity of nucleotide analogs in a primer extension assay by mitochondrial DNA-dependent RNA polymerase (22). Using a similar method, the discrimination values of nucleotide analogs, measured in the SARS-CoV-2 RdRp primer extension assay developed in this study, were employed to evaluate the relative incorporation efficiency of nucleotide analogs by SARS-CoV-2 RdRp. An initial time-course experiment showed that the incorporation of natural rNTPs by SARS-CoV-2 RdRp is very fast (data not shown) and RNA synthesis was finished within 1 min after adding rNTP (100 μM). To get a quantitative measurement of *K_1/2_* value (nucleotide concentration resulting in 50% product extension), a short reaction time (20 s) after adding different nucleotide analogs were used. Fig. 7A and B show the primer/template design and the results of testing of ATP and two ATP analogs (Remdesivir-TP and 6-Chloropurine-TP). The measured values of *K_1/2_* (the nucleotide concentration at which half of the 31-mer product is extended to the 32-mer product) for ATP (*K*_1/2, ATP_), Remdesivir-TP (*K*_1/2, Remdesivir-TP_) and 6-Chloropurine-TP (*K*_1/2, 6-Chloropurine-TP_) were 0.04167 μM, 0.03305 μM, and 3.351 μM respectively, and the calculated discrimination values for *D*_Remdesivir-TP_ and *D*_6-Chloropurine-TP_ were 0.79 and 80, respectively (Fig. 7C). This result suggested that Remdesivir-TP was incorporated into RNA by SARS-CoV-2 RdRp more efficiently than natural ATP. The incorporation efficiency of other ATP analogs by SARS-CoV-2 RdRp was much lower than that of ATP. To get a quantitative measurement of *K_1/2_* values, a longer reaction time (15 min) is needed to increase the percentage of incorporation of nucleotide analog by SARS-CoV-2 RdRp. Since the *K_1/2_* of natural ATP is impossible to measure directly under such conditions (it is below the limit of sensitivity of the assay), the *K_1/2_* value of 6-Chloropurine-TP was used as a surrogate comparator. A similar strategy has been used to evaluate the incorporation efficiency of nucleotide analog by mitochondrial DNA-dependent RNA polymerase (22). Fig. 7D-E shows the testing of ATP analogs and discrimination value calculation. In this assay, the *K_1/2_* values of several ATP analogs were measured and compared to the *K_1/2_* value of 6-Chloropurine-TP, which was used as a reference to calculate the *D*\*_ATP analog_ value (where *D*\*_ATP analog_ = *K*_1/2, ATP analog_/*K*_1/2, 6-Chloropurine-TP_). The discrimination values of the different ATP analogs are summarized in Table 1. *D^cal^* is a calculated discrimination value obtained using the equation *D^cal^* _ATP analog_ = *D*\*_ATP analog_ × *D*_6-Chloropurine_. *D^cal^* represents the discrimination of incorporation by RdRp between natural ATP and a tested ATP analog. Mis-incorporation of GTP base-pairing with uridine in the template was also measured, which can be used as a guideline to evaluate the possibility of nucleotide analogs incorporation in the cell. If the incorporation efficiency of an ATP analog is lower than GTP misincorporation efficiency, it probably has less chance to be incorporated in the cell, unless very high intracellular nucleotide analogs concentration can be reached. As shown in Table 1, *D^cal^* values for Remdesivir-TP, 6-Chloropurine-TP, Clofarabine-TP, Ribavirin-TP, Favipiravir-TP, Tenofovir-DT and GTP were 0.78 ± 0.02, 78.0 ± 3.5, >112242, 24999 ± 828, 7343 ± 752, >112242 and 8683 ± 600, respectively. Based on comparison with *D*^cal^_GTP_ (natural GTP misincorporation as ATP), Remdesivir-TP, 6-Chloropurine-TP can be incorporated into RNA very efficiently by RdRp; Ribavirin-TP and Favipiravir-TP showed less possibility to be incorporated into RNA by RdRp. Only a small fraction of primer was extended in the condition of Clofarabine-TP and Tenofovir-DP in the primer extension condition used in this assay, which suggested that the incorporation efficiency of those two nucleotides is very low. Using a similar strategy, several GTP analogs (Fig. S1) and UTP analogs (Fig. S2) were also be tested and the data were summarized in Table 1. For GTP analogs, 6-Thio-GTP and 2’-C-Methyl-GTP can be incorporated into RNA very efficiently; the incorporation efficiency of 6-Methylthio-GTP, Oxo-GTP, Remdesivir-TP, Ribavirin-TP, and Favipiravir-TP is lower than or close to the misincorporation of ATP base-pairing with cytidine in the template, which suggested that they may not be incorporated into RNA as GTP analog in the cell. For UTP analogs, 2’-Amino-UTP, 2’-Azido-UTP and ara-UTP showed higher incorporation efficiency; incorporation efficiency for Stavudine-TP, 2’-O-Methyl-UTP, Sofosbuvir-TP, and Ribavirin-TP was very low. Remdesivir-TP and Favipiravir-TP were also tested as UTP analogs and no incorporation was observed in the condition used in this assay (15 min incubation) (data not shown).

**Table 1.** Discrimination values of nucleotide analog by SARS-CoV-2 RdRp

| Nucleotide analog | $D^*_{analog}$ (mean ± SD) | $D^{cal}_{analog}$ (mean±SD) |
| --- | --- | --- |
| **ATP analogs** |  |  |
| Remdesivir -TP | | $0.78 \pm 0.02$ |
| 6-Chloropurine-TP | 1 | $78.0 \pm 3.5$ |
| Clofarabine-TP | >1439 | >112242 |
| Ribavirin-TP | $320.5 \pm 10.6$ | $24999 \pm 828$ |
| Tenofovir-DP | >1439 | >112242 |
| Favipiravir-TP | $94.1 \pm 9.6$ | $7343 \pm 752$ |
| GTP | $111.3 \pm 7.7$ | $8683 \pm 600$ |
| **GTP analogs** |  |  |
| 6-Thio-GTP | | $10.6 \pm 1.1$ |
| 2'-C-Methyl-GTP | 1 | $61.7 \pm 9.0$ |
| 6-Methylthio-GTP | $73.2 \pm 14.3$ | $4516 \pm 884$ |
| Oxo-GTP | $37.6 \pm 9.7$ | $2317 \pm 600$ |
| Remdesivir-TP | $1786.4 \pm 173.8$ | $110222 \pm 10726$ |
| Favipiravir-TP | >2319 | >143089 |
| Ribavirin-TP | >2319 | >143089 |
| ATP | $55.5 \pm 12.9$ | $3427 \pm 797$ |
| **UTP analogs** |  |  |
| 2'-Amino-UTP | 1 | $75 \pm 5$ |
| 2'-O-methyl-UTP | $102.5 \pm 5.1$ | $7688 \pm 380$ |
| 2'-Azido-UTP | $11.1 \pm 0.5$ | $831 \pm 35$ |
| ara-UTP | $8.3 \pm 0.5$ | $625 \pm 40$ |
| Stavudine-TP | $54.8 \pm 6.7$ | $4109 \pm 502$ |
| Sofosbuvir-TP | $97.4 \pm 4.9$ | $7307 \pm 368$ |
| Ribavirin-TP | $825.8 \pm 131.2$ | $61933 \pm 9843$ |
| CTP | $68.4 \pm 8.6$ | $5131 \pm 645$ |

$D^*_{ATP\ analog} = K_{1/2\ ATP\ analog} / K_{1/2,\ 6\text{-Chloropurine-TP}}; D^{cal}_{ATP\ analog} = D^*_{ATP\ analog} \times D_{6\text{-Chloropurine-TP}}$
$D^*_{GTP\ analog} = K_{1/2\ GTP\ analog} / K_{1/2,\ 2'\text{-C-Methyl-GTP}}; D^{cal}_{GTP\ analog} = D^*_{GTP\ analog} \times D_{2'\text{-C-Methyl-GTP}}$
$D^*_{UTP\ analog} = K_{1/2\ UTP\ analog} / K_{1/2,\ 2'\text{-Amino-UTP}}; D^{cal}_{UTP\ analog} = D^*_{UTP\ analog} \times D_{2'\text{-Amino-UTP}}$
Data are from two independent experiments.

**Fig. 7.**
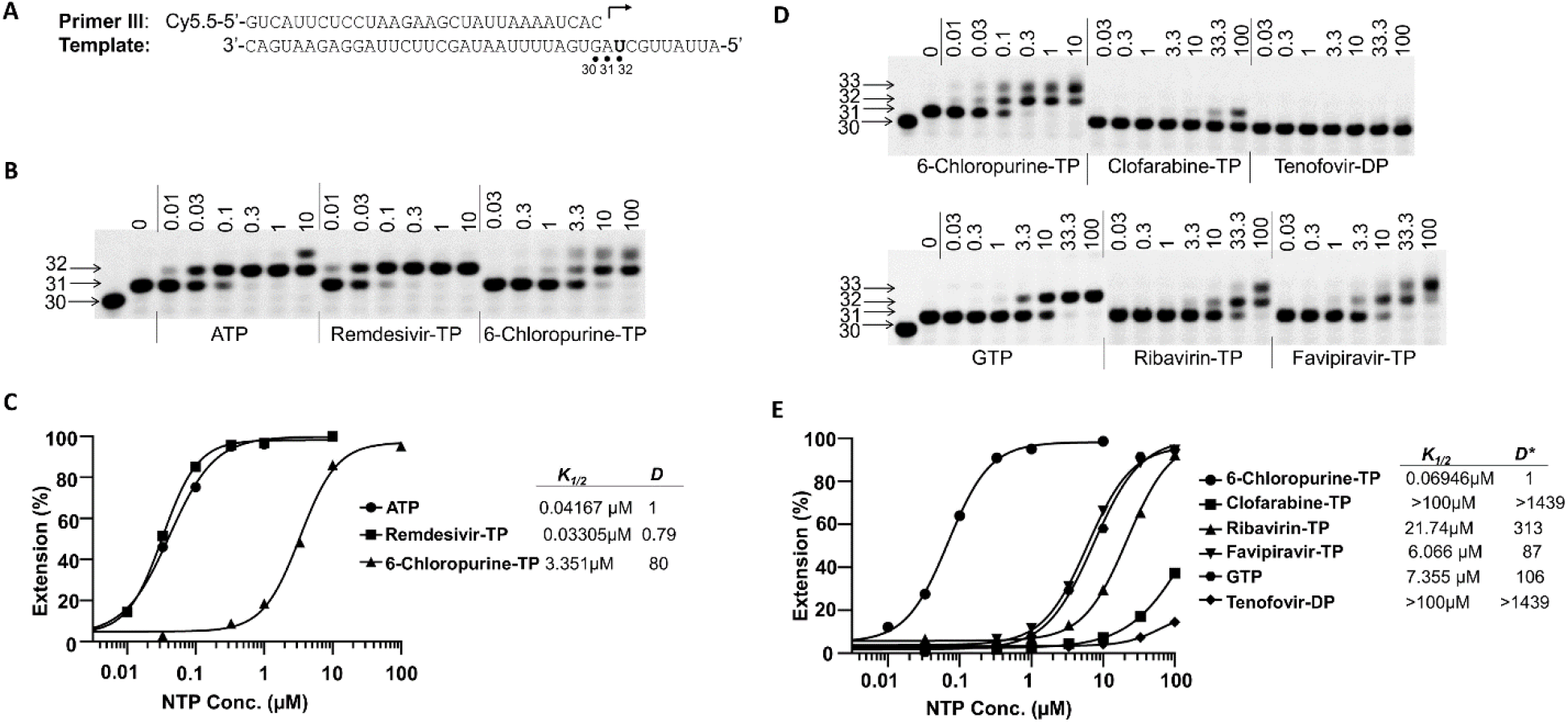
Measurement of the discrimination values of ATP analogs. **(A)** The primer and template used to assay ATP analog shown in panel B and D. **(B)** A representative image of the results of the analysis of *D*_remdesivir-TP_ and *D*_6-Chloropurine-TP_ values. Nsp12 (50 nM), and nsp8-7 (2 μM) were incubated with 5 nM P/T and 0.1 μM UTP (the first nucleotide to be incorporated) in reaction buffer for 30 min at 37 °C and then rapidly mixed with different concentrations (in micromolar) of ATP, Remdesivir-TP or 6-Chloropurine-TP, as indicated above each lane. The reactions were continued at 22 °C for 20 s before the addition of stopping solution, and the products were resolved by denaturing PAGE. The identity of the tested nucleotide is indicated at the bottom of the gel. The location of 30-mer primer, and 31-mer and 32-mer (first and second nucleotide extension products, respectively) are indicated on the left of the gel. **(C)** Quantitative analysis of ATP, Remdesivir-TP, and 6-Chloropurine-TP incorporation in the assay whose results are shown in panel B. The incorporation efficiency was evaluated based on the extension of 31-mer to 32-mer products. The measured *K_1/2_* values for ATP, Remdesivir-TP, and 6-Chloropurine-TP in this experiment were 0.04167 μM, 0.03305 μM, and 3.351 μM, respectively. The discrimination values were calculated using the equation as *K*_1/2, ATP analog_/*K*_1/2, ATP_ and are shown on the right of the graph. **(D)** A representative image of the result of the analysis of ATP analogs. Primer extension reactions were performed using a similar method described in Panel B. After adding different concentrations of ATP analog, the reactions were continued at 22 °C for 15 min before adding stopping solution. **(E)** Quantitative analysis of ATP analogs incorporation in the assay whose results are shown in panel D. *K_1/2_* analysis is as same as described in panel B and C. The discrimination between ATP analog and 6-Chloropurine-TP, *D*\*_ATP analog_, was calculated as *K*_1/2, ATP analog_/*K*_1/2, 6-Chloropurine_, and the values are shown on the right of the graph. The discrimination between ATP analogs and natural ATP, *D^cal^*_ATP analog_ was calculated as *D*\*_ATP analog_ × D_6-Chloropurine-TP_. *D*_6-Chloropurine-TP_ is 78.0 ± 3.5 (average of 2 independent experiments, one is shown in panel C). For Clofarabine-TP, Ribavirin-TP, Tenofovir-DP, Favipiravir-TP and GTP (mis-incorporated as ATP) calculated values of *D^cal^* _ATP analog_ were >112242, 24999 ± 828, >112242, 7343 ± 752 and 8683 ± 600, respectively (average of 2 independent experiments, one is shown in this figure).

### Influence of Remdesivir-TP incorporation to RNA synthesis

It was shown that Remdesivir-TP could be incorporated into RNA, and this incorporation caused delayed chain termination (23). In this report, the influence of Remdesivir-TP incorporation to the following nucleotide incorporation during RNA synthesis was tested in primer extension assay (Fig. 8). To rule out the possibility of premature RNA synthesis caused by RNA sequence variation, poly-A sequence was used downstream of uridine (Remdesivir-TP incorporation site) in the template (Fig. 8A, C, E, G). In this study, 4 different templates (shown on the top of each gel image in Fig. 8) were used to test the influence of single, double, triple, or quadruple incorporations of Remdesivir-TP to the following nucleotide incorporation. After incorporation of ATP or Remdesivir-TP, different concentrations of UTP were added to test the RNA synthesis by RdRp. Comparing with ATP, single incorporation of Remdesivir-TP did not lead to chain termination or delayed chain termination (Fig. 8B). Interestingly, the incorporation efficiency of UTP after single Remdesivir-TP incorporation was increased evidenced by a stronger 33-mer band at a low concentration of UTP in the condition of Remdesivir-TP comparing with that in the condition of ATP (Fig. 8B). Fig. 8C-D showed double incorporation of Remdesivir-TP also did not lead to chain termination or delayed chain termination. Fig. 8E-F showed triple incorporation of Remdesivir-TP did decrease the RNA synthesis efficiency after Remdesivir-TP incorporation evidenced by the 34-mer band persisting at a high concentration of UTP (Fig. 8F), which suggests a partial termination caused by triple incorporation of Remdesivir-TP. Fig. 8G-H showed quadruple incorporation of Remdesivir-TP decreased the RNA synthesis a lot (Fig. 8H, 35-mer band), which suggests a strong termination effect caused by quadruple incorporation of Remdesivir-TP.

**Fig. 8.**
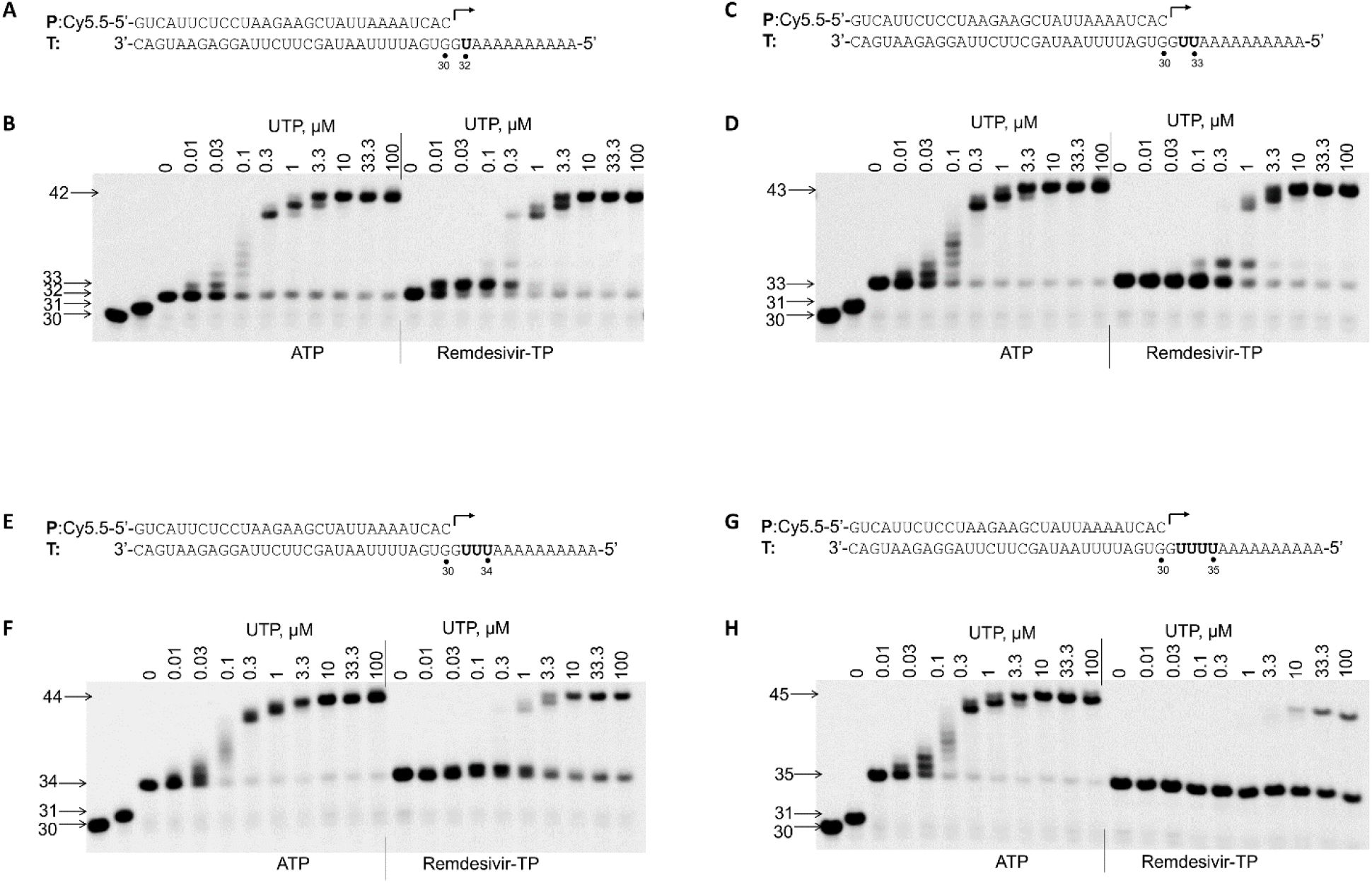
Influence of Remdesivir-TP incorporation to the following nucleotide incorporation. **(A)** The P/T used in the experiment whose result is shown in panel B. Bold “**U**” in the template indicates the position base-pairing with ATP or Remdesivir-TP. **(B)** The influence of single Remdesivir-TP incorporation to the following nucleotide incorporation. Nsp12 (50 nM), and nsp8-7 (2 μM) were incubated with 5 nM P/T (shown in panel A) and 0.1 μM CTP (the first nucleotide to be incorporated) and 0.1 μM ATP or Remdesivir-TP (the second nucleotide to be incorporated) in reaction buffer for 30 min at 37 °C and then rapidly mixed with different concentrations (in micromolar) of UTP as indicated above each lane. The reactions were continued at 22 °C for 20 s before the addition of stopping solution, and the products were resolved by denaturing PAGE. The identity of the tested nucleotide (ATP or Remdesivir-TP) is indicated at the bottom of the gel. The location of 30-mer primer and 31-mer (CTP incorporation), 32-mer (single ATP or Remdesivir-TP incorporation), 42-mer (full-length extension products), respectively, are indicated on the left of the gel. The influence of double, triple, and quadruple incorporations of Remdesivir-TP to the following nucleotide incorporation was analyzed using a similar method and shown in panel **C-D**, panel **E-F** and panel **G-H**, respectively.

## Discussion

In this study, SARS-CoV-2 nsp12, nsp8, and nsp7 were successfully constructed, expressed, and purified from *E coli*. Nsp12-nsp8-nsp7 complex (RdRp) was assembled *in vitro* and showed robust RNA synthesis activity. Using purified RdRp, we have developed an *in vitro* nonradioactive primer extension assay and have demonstrated that it can be used as a tool to identify nucleotide analog substrates which could be developed into antiviral drugs against SARS-CoV-2. The primer extension assay described in this report can also be used to develop a high through-put screen assay based on the detection of PPi released from polymerase reaction to identify non-nucleotide analog polymerase inhibitors against SARS-CoV-2 (21).

It has been shown that many nucleotide analogs currently used as antiviral drugs can be incorporated into RNA by SARS-CoV-2 RNA polymerase (27, 28), which suggests that they have the potential to be developed into antiviral drug against SARS-CoV-2. As a competitive inhibitor of RNA polymerase, a nucleotide analog must compete with natural rNTP for incorporation into viral RNA. The relative incorporation efficiency of a nucleotide analog versus natural rNTP (discrimination value) is an important criterion in evaluating the antiviral potential of nucleotide analogs. Like other studies (23), our data showed that Remdesivir-TP can be incorporated into RNA as ATP analog by SARS-CoV-2 RdRp and the incorporation efficiency is higher than natural ATP (*D*_Remdesivir-TP_ = 0.78 ± 0.02). The incorporation efficiency of Ribavirin-TP and Favipiravir-TP is very low either as ATP analog or as GTP analogs (even lower than GTP or ATP mis-incorporation), which suggested that they may not be incorporated into RNA by SARS-CoV2 RdRp in the cell unless the intracellular concentration of Ribavirin-TP and Favipiravir-TP can reach to such a high level, that they can successfully compete with natural nucleotide for incorporation. Unlike their mutagenesis effect shown in influenza virus infection, the mechanism of action for Ribavirin and Favipiravir against SARS-CoV-2 may not be mediated by RdRp and need further investigation. Sofosbuvir, an approved anti-HCV drug, has been proposed to be used for treating SARS-CoV-2. Our data showed that Sofosbuvir-TP can be incorporated into RNA by SARS-CoV-2 RdRp, but the incorporation efficiency is very low and probably it cannot compete with natural UTP for incorporation by SARS-CoV-2 RdRp in the cell. Remdesivir has been used as antiviral drug against SARS-CoV-2 and number of studies have been performed to study the antiviral mechanism of action of Remdesivir (23, 29, 30). It has been shown that incorporation of Remdesivir-TP caused delayed chain termination, which in turn block viral RNA synthesis. The influence of Remdesivir-TP incorporation to RNA synthesis catalyzed by SARS-CoV-2 RdRp was studied using primer extension assay. Our data showed that single or double incorporation of Remdesivir-TP did not lead to chain termination or delayed chain termination. Quadruple incorporation of Remdesivir-TP caused a strong termination, but no delayed chain termination was observed. There are a few parameters that may have affected the results, such as the primer/template scaffold used in different studies, template sequence downstream of Remdesivir-TP incorporation and purification, and assembling method of SARS-CoV-2 RdRp (nsp12-nsp8-nsp7 complex). Further mechanistic study is needed to address the discrepancy between our result and those in the paper of Gordon CJ et al.

## Acknowledgement

This study was supported by Bill & Melinda Gates Foundation, Tsinghua University and Beijing Municipal Government.

## Abbreviations

RdRp: RNA-dependent RNA polymerase
*D* value: Discrimination value
rNTP: ribonucleotide-5’-triphosphate
P/T: primer/template duplex

---

**Fig. S1.**
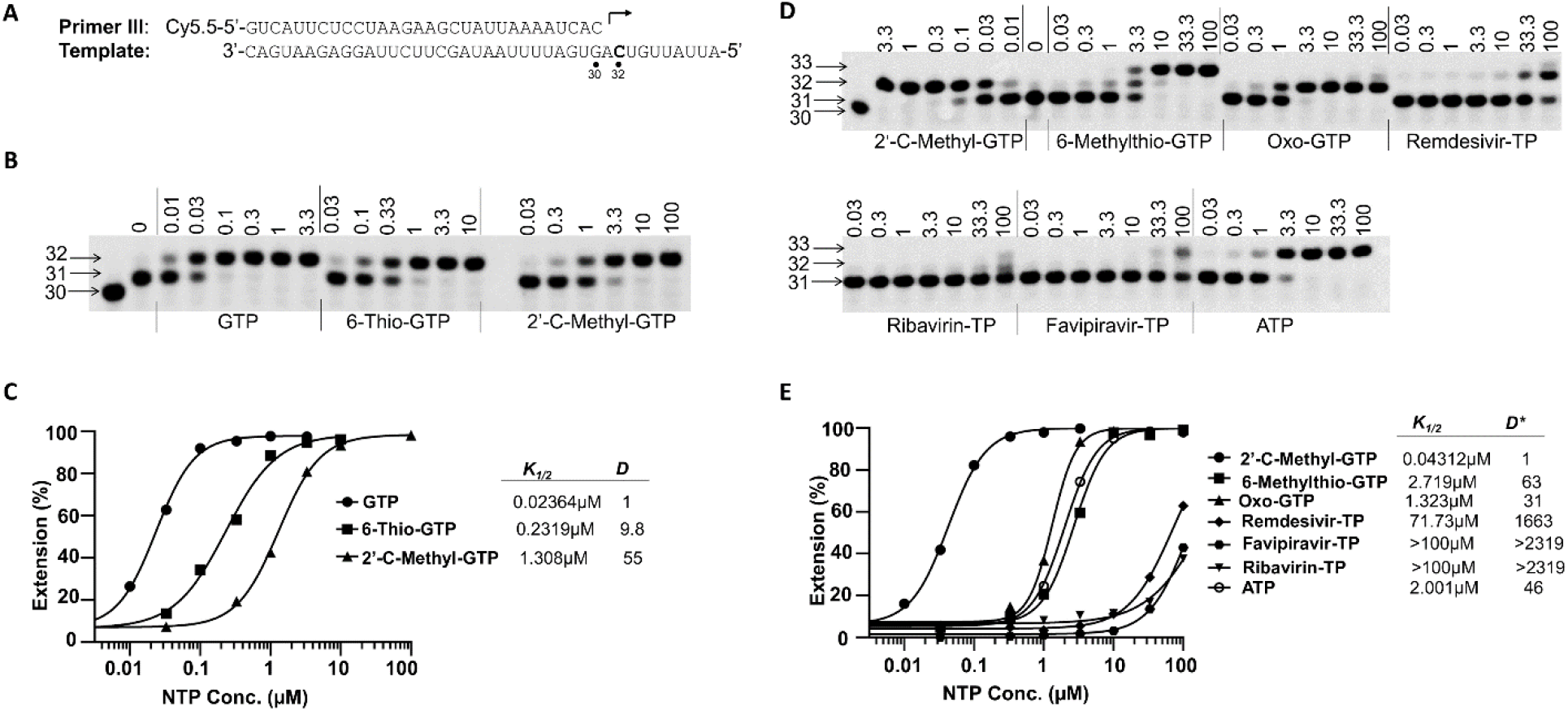
Measurement of the discrimination value of GTP analogs. **(A)** The primer and template used to assay GTP analog shown in panel B and D. **(B)** A representative image of the results of the analysis of D_6-Thio-GTP_ and D_2’-C-Methyl-GTP_ values. Nsp12 (50 nM), and nsp8-7 (2 μM) were incubated with 5 nM P/T and 0.1 μM UTP (the first nucleotide to be incorporated) in reaction buffer for 30 min at 37 °C and then rapidly mixed with different concentrations (in micromolar) of GTP, 6-Thio-GTP or 2’-C-Methyl-GTP, as indicated above each lane. The reactions were continued at 22 °C for 20 s before the addition of stopping solution, and the products were resolved by denaturing PAGE. The identity of the tested nucleotide is indicated at the bottom of the gel. The location of 30-mer primer and 31-mer and 32-mer (first and second nucleotide extension products, respectively) are indicated on the left of the gel. **(C)** Quantitative analysis of GTP, 6-Thio-GTP, and 2’-C-Methyl-GTP incorporation in the assay whose results are shown in panel B. The incorporation efficiency was evaluated based on the extension of 31-mer to 32-mer products. The measured *K_1/2_* values for GTP, 6-Thio-GTP, and 2’-C-Methyl-GTP in this experiment were 0.02364 μM, 0.2319 μM and 1.308 μM, respectively. The discrimination values were calculated using the equation as *K*_*1/2*, GTP analog_/*K*_*1/2*, GTP_ and are shown on the right of the graph. **(D)** A representative image of the result of the analysis of GTP analogs. Primer extension reactions were performed using a similar method described in Panel B. After adding different concentrations of GTP analog, the reactions were continued at 22 °C for 15 min before adding stopping solution. **(E)** Quantitative analysis of GTP analogs incorporation in the assay whose results are shown in panel D. *K_1/2_* analysis is as same as described in panel B and C. The discrimination between GTP analog and 2’-C-Methyl-GTP, *D*\*_GTP analog_, was calculated as *K*_*1/2*, GTP analog_/*K*_*1/2*, 2’-C-Methyl-GTP_, and the values are shown on the right of the graph. The discrimination between GTP analogs and natural GTP, *D^cal^*_GTP analog_ was calculated as *D*\*_GTP analog_ × *D*_2’-C-Methyl-GTP_. *D*_2’-C-Methyl-GTP_ is 61.7 ± 9.0 (average of 2 independent experiments, one is shown in panel C). For 6-Methylthio-GTP, Oxo-GTP, Remdesivir-TP, Ribavirin-TP, Favipiravir-TP and ATP (misincorporated as GTP) calculated values of *D^cal^*_analog_ were 4516 ± 884, 2317 ± 600, 110222 ± 10726, >143089, >143089 and 3427 ± 797, respectively (average of 2 independent experiments, one is shown in this figure).

**Fig. S2.**
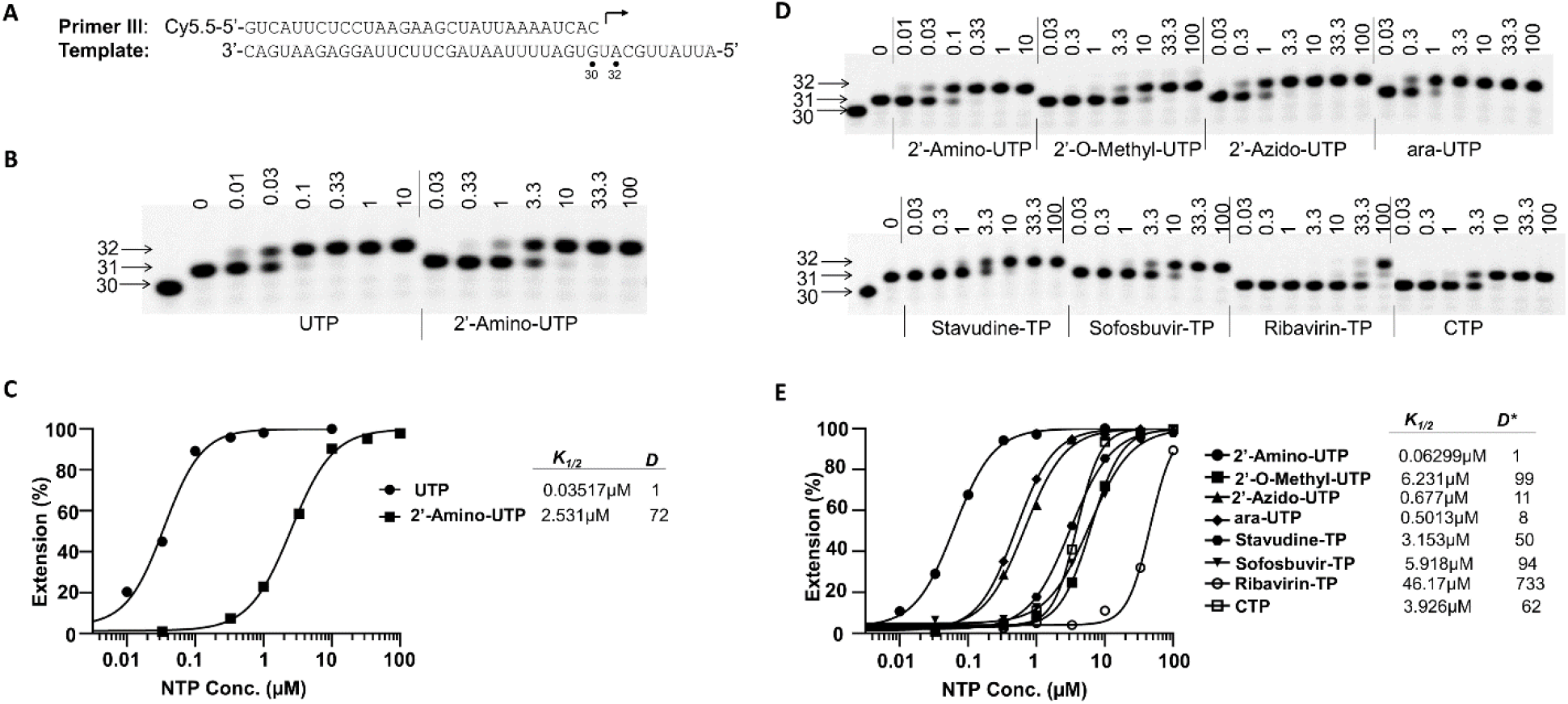
Measurement of the discrimination value of UTP analogs. **(A)** The primer and template used to assay UTP analog shown in panel B and D. **(B)** A representative image of the results of the analysis of *D*_2’-Amino-UTP_ values. Nsp12 (50 nM), and nsp8-7 (2 μM) were incubated with 5 nM P/T and 0.1 μM ATP (the first nucleotide to be incorporated) in reaction buffer for 30 min at 37 °C and then rapidly mixed with different concentrations (in micromolar) of UTP, 2’-Amino-UTP, as indicated above each lane. The reactions were continued at 22 °C for 20 s before the addition of stopping solution, and the products were resolved by denaturing PAGE. The identity of the tested nucleotide is indicated at the bottom of the gel. The location of 30-mer primer, and 31-mer and 32-mer (first and second nucleotide extension products, respectively) are indicated on the left of the gel. **(C)** Quantitative analysis of UTP, 2’-Amino-UTP incorporation in the assay whose results are shown in panel B. The incorporation efficiency was evaluated based on the extension of 31-mer to 32-mer products. The measured *K_1/2_* values for UTP, 2’-Amino-UTP in this experiment were 0.03517 μM and 2.531 μM, respectively. The discrimination values were calculated using the equation as *K*_*1/2*, UTP analog_/*K*_*1/2*, UTP_ and are shown on the right of the graph. **(D)** A representative image of the result of the analysis of UTP analogs. Primer extension reactions were performed using a similar method described in Panel B. After adding different concentrations of UTP analog, the reactions were continued at 22 °C for 15 min before adding stopping solution. **(E)** Quantitative analysis of UTP analogs incorporation in the assay whose results are shown in panel D. *K_1/2_* analysis is as same as described in panel B and C. The discrimination between UTP analog and 2’-Amino-UTP, *D*\*_UTP analog_, was calculated as *K*_1/2, UTP analog_/*K*_1/2, 2’-Amino-UTP_, and the values are shown on the right of the graph. The discrimination between UTP analogs and natural UTP, *D*^*ca*l^_UTP analog_ was calculated as *D*\*_UTP analog_ × *D*_2’-Amino-UTP_. *D*_2’-Amino-UTP_ is 75 ± 5 (average of 2 independent experiments, one is shown in panel C). For 2’-O-Methyl-UTP, 2’-Azido-UTP, ara-UTP, Stavudine-TP, Sofosbuvir-TP, Ribavirin-TP and CTP (mis-incorporated as UTP) calculated values of *D^cal^* _UTP analog_ were 7688 ± 380, 831 ± 35, 625 ± 40, 4109 ± 502, 7307 ± 368, 61933 ± 9843 and 5131 ± 645, respectively (average of 2 independent experiments, one is shown in this figure).

